# From Face to Person: Disentangling the Neural Origins of the N170, N250, and SFE in Familiar Face Recognition

**DOI:** 10.64898/2026.05.04.722329

**Authors:** Meijiao Li, Chenglin Li

**Affiliations:** School of Psychology, Zhejiang Normal University, Jinhua, China

**Keywords:** EEG, visual exposure, identity representation, person-related semantic knowledge, familiar face recognition

## Abstract

Recognizing familiar faces is essential in our everyday life. ERP studies have identified three components sensitive to face familiarity (N170, N250, and SFE), but whether these signals arise from visual experience, identity information, or semantic knowledge remains to be directly tested. Using a sequential familiarization paradigm, we progressively trained the same initially unfamiliar faces with visual exposure, identity associations, and biographical knowledge, recording EEG after each phase. The N250 emerged immediately after visual familiarization and remained stable thereafter; the N170 appeared only after identity familiarization; and the SFE exhibited a graded, enhanced pattern: absent after visual exposure, emerging after identity training, and reaching maximum effect after semantic familiarization. These findings provide the first direct evidence that these three ERP markers are differentially driven by distinct types of information, revealing the temporal dynamics through which person-related knowledge transforms a face percept into the recognition of a known person.

## 1. Introduction

The ability to recognize familiar faces is essential for human social interaction, enabling us to navigate complex social environments with remarkable efficiency (Bruce & Young, 1986; Young & Burton, 2017). The cognitive and neural distinction between processing familiar and unfamiliar faces is profound: whereas unfamiliar-face perception relies heavily on image-specific pictorial cues, familiar-face recognition reflects the activation of a robust, image-invariant person representation (Burton et al., 2011; Jenkins et al., 2011). Decades of event-related potential (ERP) studies have identified a sequence of neural markers sensitive to this face familiarity distinction, most notably the N170 (Bentin et al., 1996; Caharel & Rossion, 2021), N250 (Schweinberger & Neumann, 2016), and the sustained familiarity effect (SFE; Wiese et al., 2019a, 2024). Despite the well-documented sensitivity of these components to facial familiarity (Huang et al., 2017), a fundamental question remains unresolved: What specific type of information drives each of these distinct neural markers? In other words, the precise correspondence between the accumulation of person-related information and the emergence of specific ERP familiarity markers is unknown. Resolving this issue is critical for understanding the neural origins of these components and for understanding the cognitive pathway from face perception to person recognition. The transformation of a face from unfamiliar to familiar is a gradual, multi-stage process of information accumulation (Ambrus et al., 2021). At its most basic level, repeated visual exposure alone can generate perceptual familiarity. Behavioral studies indicate that mere visual repetition can improve face discrimination (Andrews et al., 2015), and neuroimaging evidence shows that visual familiarization modulates neural activity in core face-processing regions such as the fusiform gyrus (Gobbini & Haxby, 2006; Kosaka et al., 2003). Critically, ERP studies have linked this accumulation of visual experience to the N250 component (Tanaka et al., 2006). The N250 familiarity effect refers to a more negative component for familiar relative to unfamiliar faces at occipitotemporal electrodes beginning around 200 ms, which is thought to index the successful matching of a perceptual input to a stored, domain-selective facial representation (Andrews et al., 2017; Schweinberger & Neumann, 2016). Importantly, this effect is robustly observed after even brief visual familiarization (Popova & Wiese, 2023a, 2023b), suggesting that the N250 serve as the earliest electrophysiological signature of forming visual memory traces for a face.

However, person recognition extends beyond visual memory to encompass identity-specific associations. Assigning a unique identity label (i.e. a name) to a face represents a crucial conceptual anchor (Schwartz & Yovel, 2016, 2019). Critically, recent evidence challenges the classic view that face familiarity effects begin only at the N250, suggesting instead that identity learning can modulate even earlier perceptual stages. Specifically, the N170, a face-sensitive component peaking around 170 ms and traditionally associated with structural encoding of faces (Bentin et al., 1996; Eimer, 2011), has been shown to differentiate highly familiar from unfamiliar faces when robust personal representations have been formed (for reviews, see Caharel & Rossion, 2021, but Wiese et al., 2024). In addition, recent studies using multivariate pattern analysis have revealed that identity-specific information can be decoded from neural activity within the N170 time-range (Kovács et al., 2023). These findings suggest that the sensitivity of N170 to familiar faces may reflect the consolidation of an individualized person representation that facilitates top-down perceptual tuning rather than mere face detection.

Importantly, the integration of rich, person-related semantic knowledge, including biographical details such as age, occupation, hobbies, and personality traits, contributes to familiar face recognition (Li et al., 2022; Shi et al., 2026; Young & Burton, 2017). Previous studies have shown that associating faces with such semantic information elevates from identification recognition to a deeper, more person-based knowing, substantially improving memory performance (Herzmann & Sommer, 2010; Schwartz & Yovel, 2016, 2019). In the ERP domain, this stage is most clearly associated with the SFE, a sustained negativity maximal between 400 and 600 ms at occipito-temporal sites (Wiese et al., 2019a). The SFE is substantially larger than the N250, modulated by the depth of personal relevance, and is theorized to reflect the integration of perceptual face representations with domain-general affective and semantic knowledge, likely via reentrant feedback from memory and emotion-related information (Bojdo et al., 2025; Wiese et al., 2024). Notably, the SFE is absent for faces that are only visually familiar and emerges only when a face is known as a familiar person (Wiese et al., 2019a, 2019b, 2023; Popova & Wiese, 2023a, 2023b).

Although previous studies have independently linked visual, identity, and semantic knowledge to above-mentioned neural markers, no study has systematically manipulated the type of information available for the same set of faces within a unified experimental framework. Consequently, it remains unknown whether the N170, N250, and SFE are causally driven by distinct types of information. Addressing this gap is essential for determining the cognitive origins of these ERP components during face familiarization.

To directly address this gap, the present study employed a novel sequential familiarization paradigm in which the same initially unfamiliar faces were progressively trained with three types of information. In the present study, participants were familiarized with training identities through (1) visual perceptual exposure, (2) face-identity (name) association, and (3) person-related semantic knowledge (biographical). EEG was recorded after each phase while participants performed an old/new recognition task, allowing us to isolate the unique contribution of each type of information to the emergence of the N170, N250, and SFE. We hypothesized that if the N250 primarily indexes visual memory accumulation, it should emerge immediately following the visual familiarization and show no further enhancement with subsequent. In contrast, we predicted that N170 differentiation would be contingent upon identity formation and would thus appear after face-name association. Finally, we hypothesized that the SFE would be absent after visual familiarization but would emerge and scale in magnitude with the successive learning of identity and, critically, semantic knowledge. By tracking the incremental genesis of these neural signals, the present study aims to reveal how the accumulation of person-related knowledge modulates the temporal dynamics that transform a perceptual face representation into the recognition of a fully known person.

## Research Transparency Statement

### General disclosures

#### Conflicts of interest

The authors have declared no conflicts of interest.

#### Funding

This study was supported by the Zhejiang Provincial Natural Science Foundation of China (Grant No. LQN25C090005) and the Major Humanities and Social Sciences Research Project in Zhejiang Higher Education Institutions (Grant No. 2024GH087).

#### Artificial intelligence

No artificial intelligence technologies were used in the research or writing of this article.

#### Ethics

The current study was approved by the ethics committee of Zhejiang Normal University (ZSRT2024235).

## 2. Method

### 2.1. Participants

The required sample size was determined via a priori power analysis based on the N250 and SFE difference between familiar and unfamiliar faces reported by Wiese et al (2019a; paired-sample *t* test, two-sided*, d*_z_ = .8, power = .95). Using G*Power 3.1.9.7 (Faul et al., 2007), this analysis suggested a minimum sample size of N = 23. To enhance the power further and account for potential attrition during the training, thirty-one participants were initially recruited and received monetary compensation for their participation. Six participants were excluded from the final analysis for following reasons: one for excessive anticipatory responses (17.3% of all trials); one for failure to follow task instructions during one EEG session (*d’* = -0.192); two for a strong bias toward one response key; and two for very low accuracy (< 60%) in the final test, indicating failure to acquire semantic knowledge. The remaining 25 participants (16 females; mean age 21.2 ± 2.53 years) were right-handed and had normal or corrected-to-normal vision. They gave written informed consent before the experiment. The current study was approved by the ethics committee of Zhejiang Normal University and conducted in accordance with the guidelines of the Declaration of Helsinki.

### 2.2. Experimental Materials

To avoid the influence of prior experience and to ensure that the trained faces belonged to the same racial group as the participants, 24 Southeast Asian celebrities were initially selected as candidate identities. Ten additional participants were recruited to rate their familiarity with these individuals. Based on their feedback, eight unfamiliar celebrities (four males and four females) were selected. Four of these identities were assigned to the training identities, while the remaining four served as unfamiliar control identities (see Figure 1).

**Figure 1.**
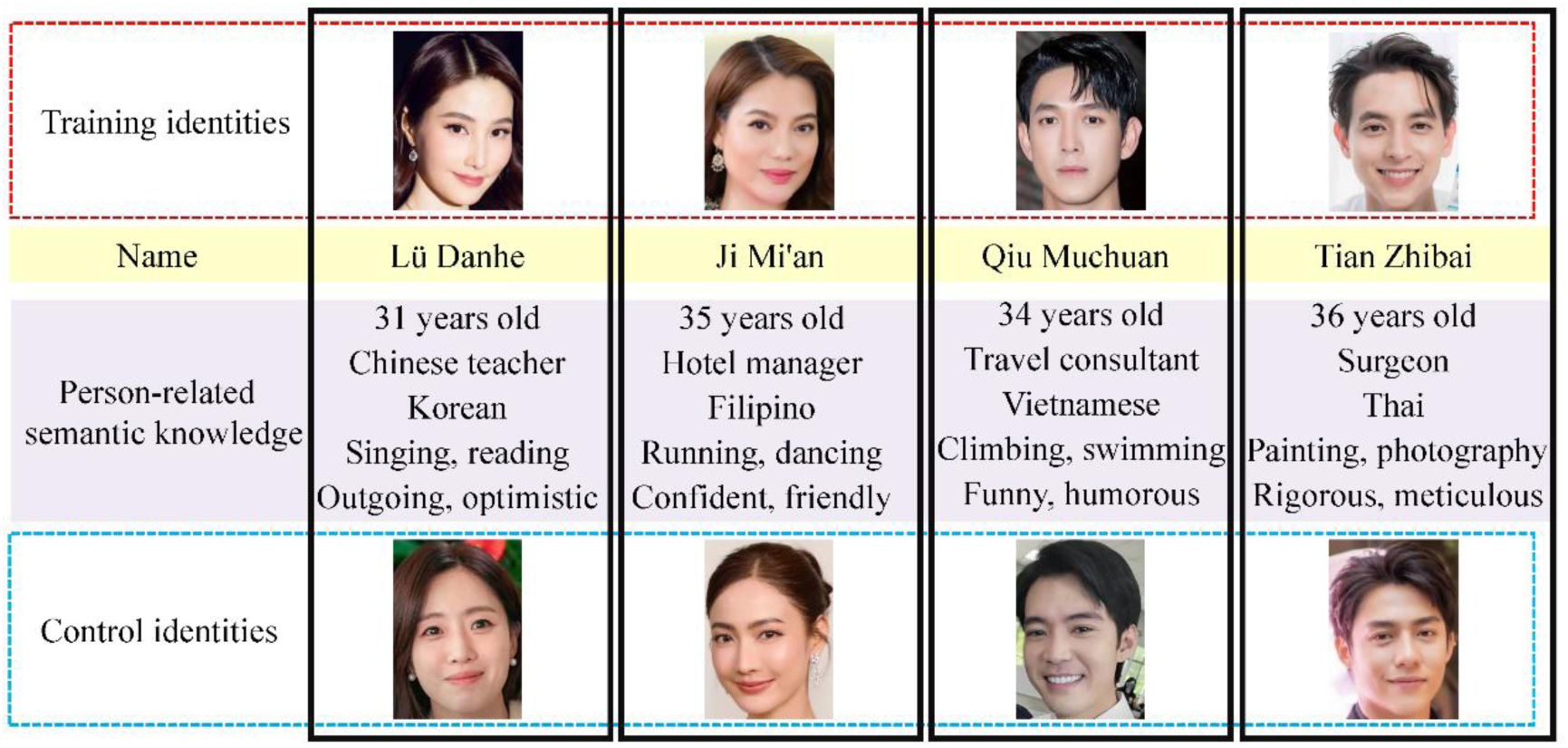
Example of Experimental Materials in Three Familiarization Phases and EEG Recording Sessions. *Note.* A sample face image of each training identity is presented, together with the artificial name and the person-related semantic knowledge. Control identities were only used during the EEG Recording Sessions.

For each of the eight selected identities, 20 color and ambient photographs were collected from the internet. All images varied naturally in viewpoint, lighting, and expression (Li et al., 2022). We used a deep neural network (DNN; Shi et al., 2026) to evaluate facial similarity between the trained identities and control identities and no significant physical differences existed between the two sets of stimuli. Faces were cropped from the original images and resized to subtend approximately 3.9° × 2.8° of visual angle at a viewing distance of 70 cm.

To minimize the influence of prior name familiarity, four artificially constructed Chinese names were used for the trained identities: 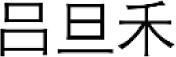 (L ü Danhe), 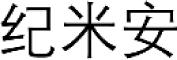 (Ji Mi’an), 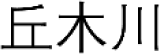 (Qiu Muchuan), 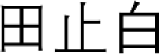 and (Tian Zhibai). These names were constructed based on the character frequency list of the Modern Chinese frequency dictionary (Wang et al., 1986), with each character having low-frequency (M = .24/1000, range: .01/1000 - .91/1000), and the number of strokes varying between 3 and 6. Three characters were randomly combined to form names that were rarely used in daily life. In addition, to increase task difficulty during name learning, three foil names were additionally generated for each identity by replacing one character of target name, and were visually and phonologically similar to the target names.

For the four trained identities, person-related semantic knowledge consisted of five categories of fictional biographical information: ages, nationality, occupations, hobbies and personality traits (Figure 1; Kovács, 2020; Li et al., 2022; Shi et al., 2026). All experimental materials are publicly available on the Open Science Framework platform (https://osf.io/e8vp9).

### 2.3. Procedure

The experiment consisted of three consecutive days, each containing a familiarization phase and a subsequent EEG recording session (Figure 2). To gradually establish person representation from perceptual exposure to identity-specific association, and to person-related semantic knowledge (Ambrus et al., 2021; Kovács, 2020), the familiarization phase was arranged in the order of visual familiarization, identity familiarization, and semantic familiarization. An EEG recording session was conducted immediately after the completion of each familiarization phase to track the contribution of different information to the neural dynamics of face familiarization. All tasks were presented on a LCD monitor (24 inch, 1920 × 1080 pixel resolution, refresh rate 60 Hz) at a viewing distance of approximately 70 cm, and the experimental program was implemented in MATLAB (R2016a) using the Psychtoolbox-3 extension.

**Figure 2.**
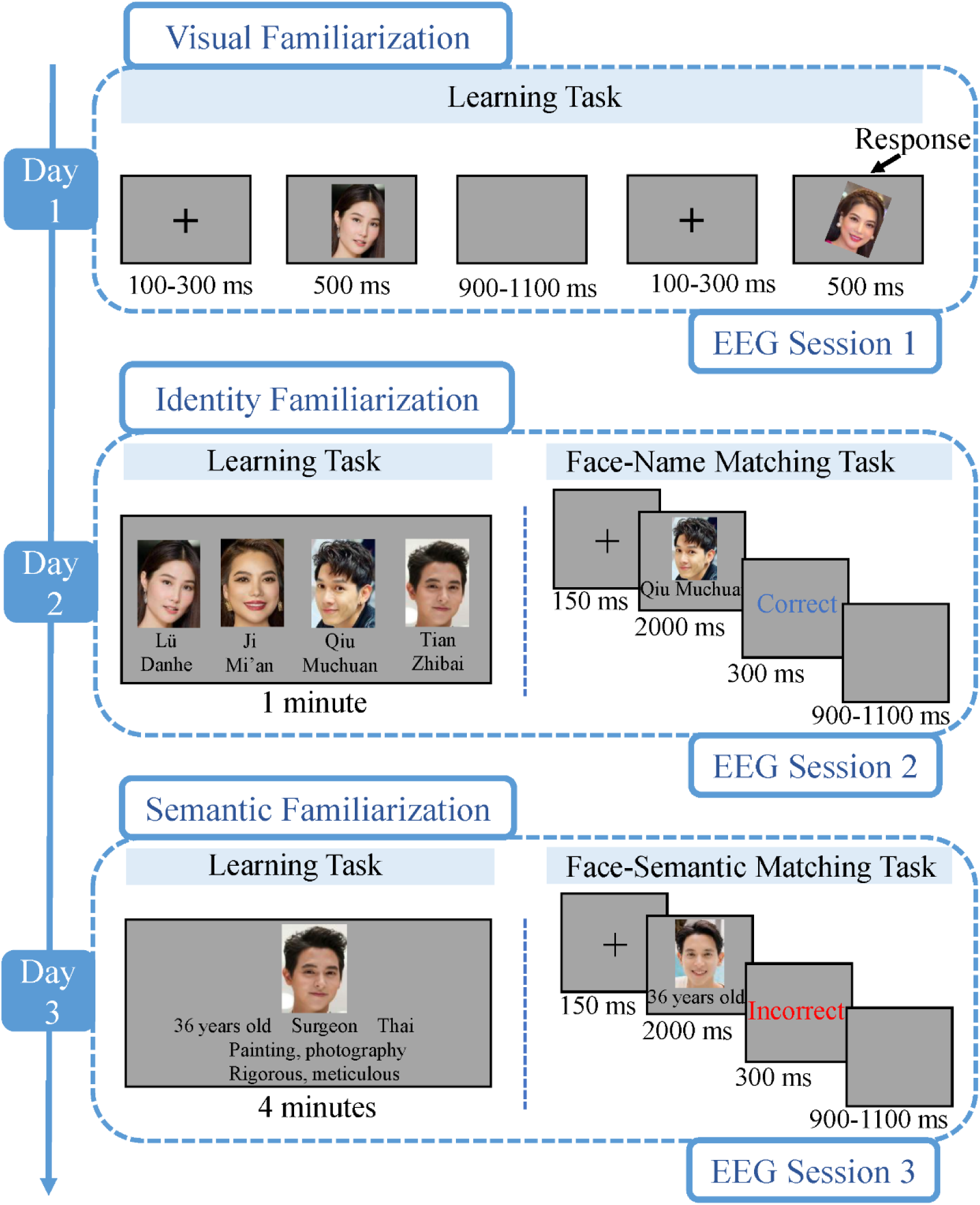
Experimental Procedure Over Three Consecutive Days. *Note.* All textual materials are presented in English, but during the experiment, the textual materials were all in Chinese.

#### 2.3.1 Familiarization Procedure

##### 2.3.1.1. Visual Familiarization

To establish perceptual familiarity with trained identities through repeated visual exposure (Andrews et al., 2015; Olivares et al., 1994), four of the eight identities were randomly selected as training identities (two males and two females). For each of these four identities, five ambient images of each training identity were selected for the learning task.

The learning task was designed as an oddball paradigm consisting of 4 blocks of 264 trials in total (240 non-target trials and 24 target trials). On non-target trials, a face image was presented centrally, and participants were instructed to attend to the images carefully. On target trials, the face image was tilted 10° clockwise or counterclockwise, and participants were required to press the space button as quickly as possible. Each trial began with a fixation cross presented for 100-300 ms, followed by the face stimulus for 500 ms. The inter-trial-interval (ITI) ranged from 900 to1100 ms.

##### 2.3.1.2. Identity Familiarization

Following visual familiarization, the same four identities were carried forward to the identity familiarization phase, during which they were associated with the artificially constructed, low-frequency names. The face images remained identical to those used in the visual familiarization phase. The training procedure was divided into a learning task and a face-name matching task. In the learning task, the faces of the four identities were displayed simultaneously on the screen, with the corresponding name printed below each face. Participants were given 1 minute to memorize these face-name pairings.

To enhance learning and minimize reliance on low-level cues, we used three foil names as distractors in the face-name matching task, which consisted of 4 blocks with 160 trials in total. Each trial started with a fixation cross for 150 ms, followed by the face and name display for 2000 ms. Participants were asked to judge whether the name matched the face. Immediate accuracy feedback was presented for 300 ms, followed by a variable ITI of 900-1000 ms. The training continued until participants achieved an overall accuracy of at least 90%, ensuring robust acquisition of face-name associations. On average, participants completed this task in 13 minutes.

##### 2.3.1.3. Semantic Familiarization

After identity familiarization, the same four identities were further associated with person-related semantic knowledge. The face images remained unchanged from the previous phases. Each identity was assigned five categories of biographical information (age, nationality, occupation, hobbies, and personality traits (adapted from Shi et al., 2026). During the learning task, participants were given a total of 4 minutes to memorize the face-semantic information correspondences. This was followed by a face-semantic matching task, where face and semantic information were presented, and participants were asked to judge whether the semantic information matched the face. The trial structure and timing parameters (fixation: 150 ms; stimulus: 2000 ms; feedback: 300 ms; ITI: 900-1000 ms) were identical to those used in the identity familiarization phase. The training was concluded once an overall accuracy criterion of 90% was met. On average, participants completed this phase in 19 minutes.

#### 2.3.2. EEG Recording Sessions

Each of the three familiarization phases was immediately followed by an EEG recording session. These sessions were designed to investigate the neural correlates of familiarity as different types of information were accumulated across training stages.

In each session, the stimulus set consisted of five previously unseen ambient images of each trained identity and five images of each unfamiliar control identity, ensuring that any observed effects reflected identity familiarity rather than image-specific repetition. All sessions employed an old/new recognition task, but the definition of “old” was contingent upon the specific familiarization stage. For example, *Session 1 (Post-Visual Familiarization)*: Participants were instructed to judge whether the presented face was “old” (i.e., visually exposed during the oddball task) or “new” (i.e., completely novel control faces). *Session 2 (Post-Identity Familiarization)*: Participants were instructed to judge whether the presented face was “old” (i.e., associated with a learned name) or “new” (i.e., faces without learned name information). *Session 3 (Post-Semantic Familiarization)*: Participants were instructed to judge whether the presented face was “old” (i.e., associated with learned person-related semantic knowledge) or “new” (i.e., faces without any learned semantic information).

Each session consisted of 8 blocks of 320 trials in total. The trial structure was identical across sessions: a fixation cross was presented for 500 ms, followed by a face image for 1000 ms. After a brief blank interval of 200 ms, a “Response” cue appeared on the screen, at which point participants were asked to complete the old/new recognition task. The ITI varied randomly between 900 and 1100 ms.

### 2.4. EEG Recording and Preprocessing

The electroencephalogram (EEG) was recorded continuously from 64 Ag/AgCl scalp electrodes according to the International 10-20 system. Two Brain Amp DC amplifiers (Brain Products GmbH, Germany) were used for signal acquisition. The electrode AFz served as the ground, and TP9 was used as the recording reference. Electrode impedances were kept under 5kΩ during the recording sessions. The EEG signal was digitized at a sampling rate of 1000 Hz, and no online filtering was implemented during acquisition.

Offline preprocessing was conducted using the EEGLAB (version 2025.0.0) and ERPLAB (version 12.01). Electrode localizations were aligned to the BESA template, and some channels identified as bad channels during visual inspection were interpolated using spherical spline interpolation. After removing electrooculogram (EOG) channels, the data were re-referenced to the common average of all remaining scalp electrodes. Following the preprocessing pipeline of Wiese et al. (2019a), the continuous EEG was band-pass filtered between 0.1 and 40 Hz using a finite impulse response (FIR) filter, and a notch filter was applied between 49 and 51 Hz to attenuate line noise. Artifact correction was performed using independent component analysis (ICA) to identify and remove components associated with ocular and muscular artifacts. Subsequently, only trials with correct behavioral responses were retained for further analysis. The data were segmented into epochs extending from -200 to 1000 ms relative to stimulus onset and baseline-corrected using the 200 ms pre-stimulus interval. An automatic artifact rejection procedure was applied to exclude any epoch in which the absolute voltage exceeded ±100 μV at any scalp electrode. The proportion of rejected epochs did not exceed 10% in any condition.

### 2.5. Behavioral and EEG Data Analysis

#### 2.5.1. Behavioral Performance

To test the familiarization effect, we analyzed the accuracy and reaction times of the old/new recognition task in the EEG session. In addition, detection sensitivity (*d’*) and response bias (criterion C) were further calculated for each participant based on their performance in the old/new recognition task. Firstly, hit rates were calculated from trials in which an “old” (training) identity was correctly recognized, and false alarm rates were calculated from trials in which a “new” (control) identity was incorrectly judged as “old”. Then, *d’* and C were calculated based on the signal detection theory (Tanner & Swets, 1954). Trials with reaction times exceeding 3 standard deviations (SD) above each participant’s mean reaction time were excluded from the analysis. On average, 1.50% of trials were excluded per participant. The mean number of valid trials per condition was 155 (SD = 3.76).

#### 2.5.2. ERP analysis

ERP analyses focused on components previously reported in which are related to face familiarity and identity representation. Following prior work (Li et al., 2019; Wiese et al., 2019b, 2024), the peak amplitudes of the N170 were quantified between 130 and 220 ms at the lateral occipitotemporal electrodes P7 and P8. For the N250, mean amplitudes were extracted from electrodes TP9 and TP10 within the time window of 200-400 ms. Additionally, the SFE, a prolonged posterior negativity associated with high face familiarity, was assessed by measuring the mean amplitudes between 400 and 600 ms at the same TP9/TP10 electrode sites. All ERP components were conducted repeated-measures analyses of variance (ANOVAs) with the within-subjects factors Hemisphere (left, right), Identity (training, control), and Familiarization Stage (visual, identity, semantic). Greenhouse-Geisser corrections were applied when the assumption of sphericity was violated, and all multiple comparisons of post-hoc tests were corrected with the Bonferroni’s method. All code and data can be accessed on the Open Science Framework platform (https://osf.io/e8vp9).

## 3. Results

### 3.1. Behavioral Results

#### 3.1.1. Accuracy and Reaction Times

One-way repeated-measures ANOVAs were conducted on accuracy (ACC) and reaction times (RTs), with Familiarization Stage (visual, identity, semantic) as the within-subjects factor (see Figure 3).

**Figure 3.**
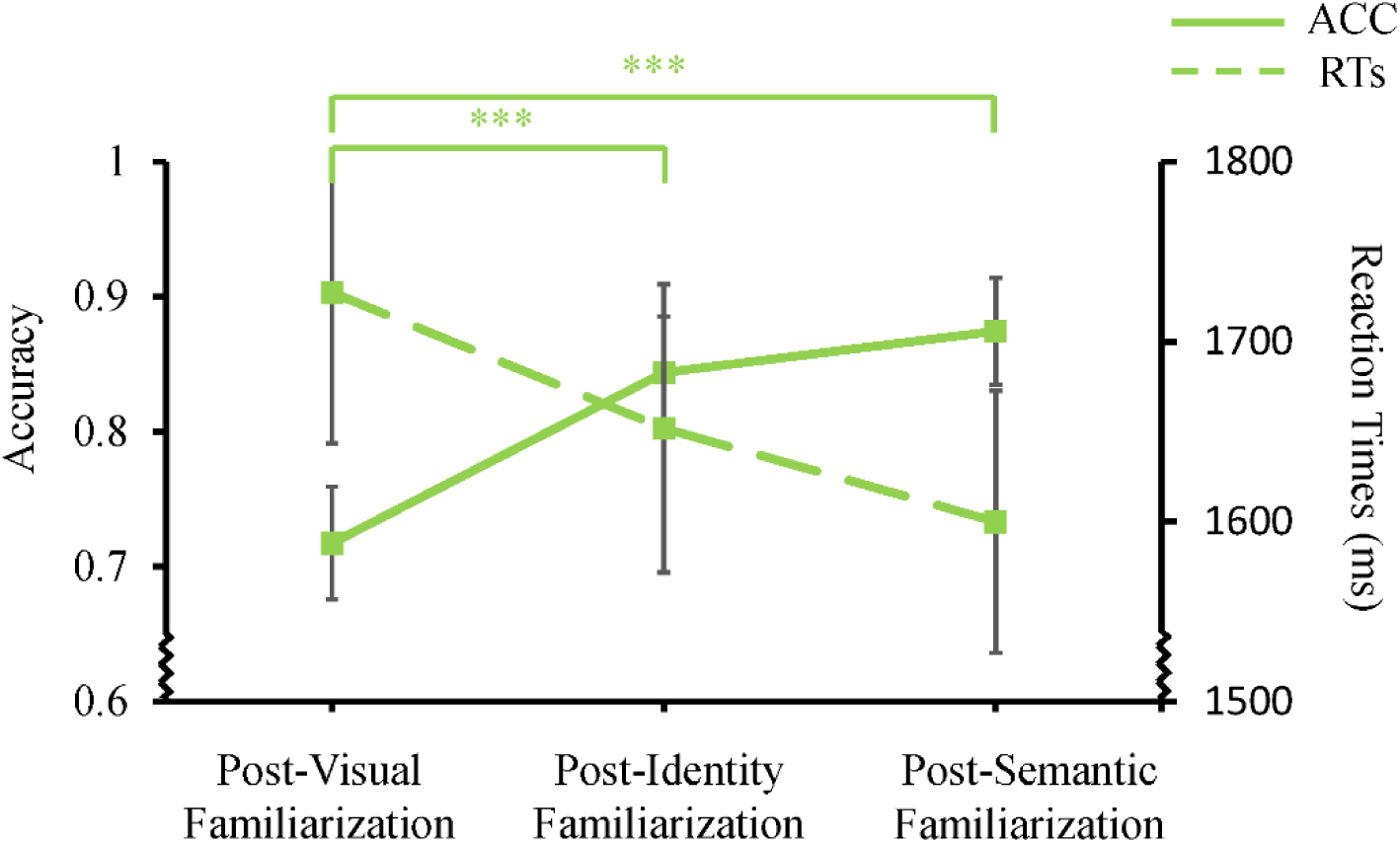
Results of Accuracy and Reaction Times in Three Old/New Recognition Tasks. *Note.* Error bars for the means indicate 95% CIs. \*\*\**p* < .001.

For ACC, the main effect of Familiarization Stage was significant (*F*_(2, 48)_ = 32.53, *p* < .001, η *^2^* = .575). Post hoc pairwise comparisons revealed that ACC was significantly lower following visual familiarization compared to both identity familiarization (mean difference = .13, 95% CI = [.07, .19]) and semantic familiarization (mean difference = .16, 95% CI = [.10, .22]). No significant difference was observed between identity and semantic familiarization (mean difference = .03, 95% CI = [.01, .07]). Thus, both identity and semantic information effectively improve the accuracy of face recognition during familiarization relative to visual exposure alone.

For RTs, the main effect of Familiarization Stage was significant (*F*_(2, 48)_ = 12.73, *p* < .001, η*_p_^2^* = .347), post hoc pairwise comparisons revealed that RT was significantly longer following visual familiarization compared to both identity familiarization (mean difference = 76 ms, 95% CI = [12, 139]) and semantic familiarization (mean difference = 127 ms, 95% CI = [50, 205]). No significant difference was observed between identity and semantic familiarization, mean difference = 52 ms, 95% CI = [-2, 106]. The decreased in RTs across familiarization stages suggests that the addition of identity and semantic knowledge facilitated faster recognition responses.

#### 3.1.2. Detection Sensitivity (*d’*) and Response Bias (C)

To evaluate how recognition sensitivity changed across familiarization stages, one-way repeated-measures ANOVAs were conducted on *d’* and C, with Familiarization Stage (visual, identity, semantic) as the within-subjects factor. The descriptive results are shown in Figure 4.

**Figure 4.**
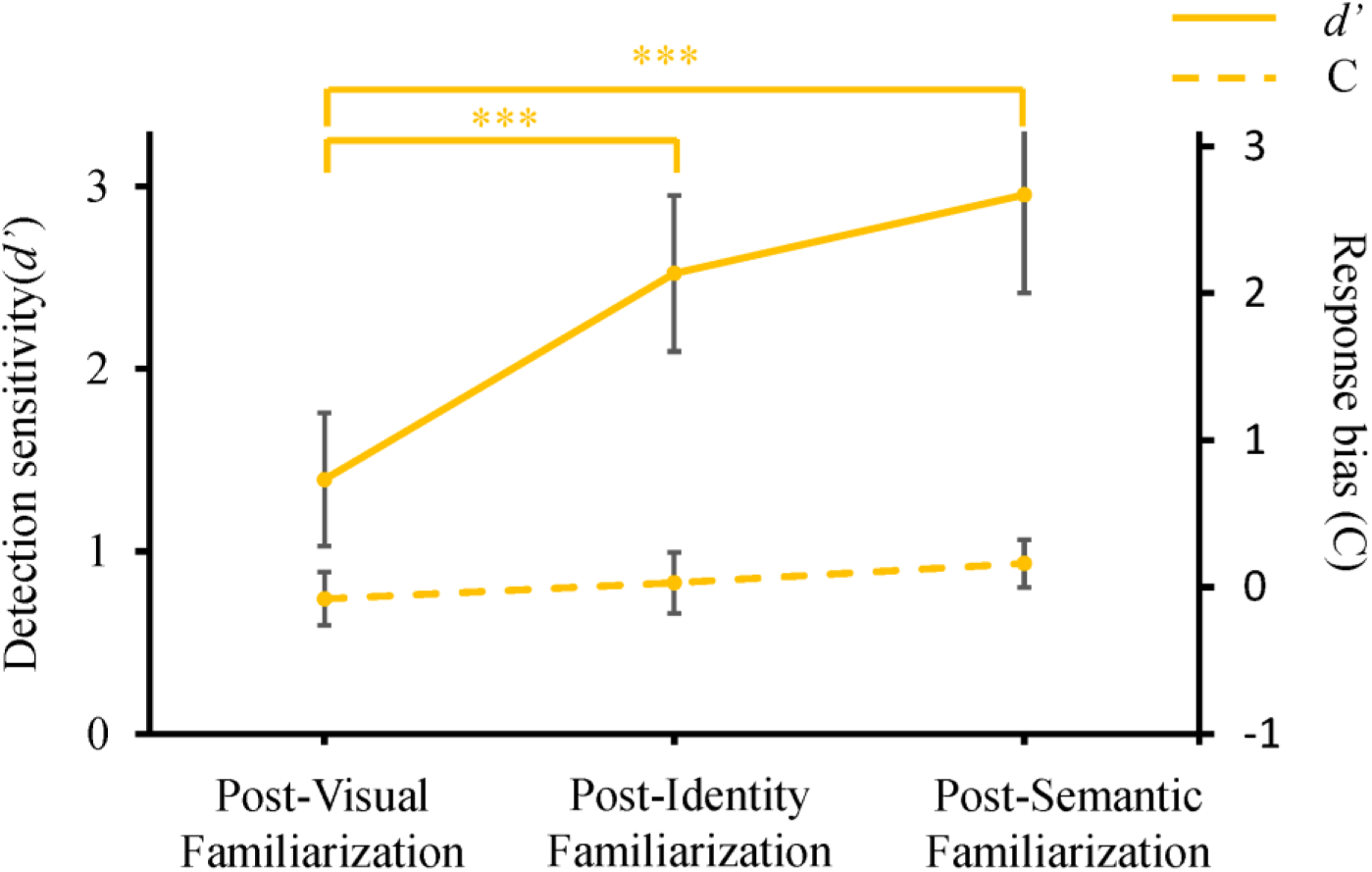
Results of Detection Sensitivity (d’) and Response Bias (C) in Three Old/New Recognition Tasks. *Note.* Error bars for the means indicate 95% CIs. \*\*\**p* < .001.

For *d’*, the main effect of Familiarization Stage was significant (*F*_(2, 48)_ = 31.77, *p* < .001, η*_p_^2^* = .570), post hoc pairwise comparisons revealed that *d’* was significantly lower following visual familiarization compared to both identity familiarization (mean difference = 1.13, 95% CI = [.67, 1.59]) and semantic familiarization (mean difference = 1.56, 95% CI = [.95, 2.17]). No significant difference was observed between identity and semantic familiarization (mean difference = .43, 95% CI = [-.91, .05]). Thus, recognition sensitivity improved substantially following the addition of identity and semantic information relative to visual familiarization alone.

For C, the main effect of Familiarization Stage failed to reach significance, (*F*_(2, 48)_ = 2.762, *p* = .073, η *^2^* = .103). Thus, response bias remained stable across stages, indicating that the familiarization procedure did not systematically alter participants’ response strategy.

### 3.2. ERP results

To examine the temporal dynamics of face familiarity representation with the accumulation of different types of person-related information, repeated-measures ANOVAs were performed on the mean amplitudes (or peak measures, where applicable) of the N170, N250, and the sustained familiarity effect (SFE), The analyses included the within-subjects factors Identity (training, control), Familiarization Stage (visual, identity, semantic), and Hemisphere (left, right).

#### 3.2.1. N170

The peak amplitudes of the N170 are shown in Table 1 and Figure 5. A repeated-measures ANOVA revealed that the main effect of Hemisphere was significant (*F*_(1, 24)_ = 4.549, *p* = .043, η *^2^* = .159, mean difference = 1.50 μV, 95% CI = [.05, 2.95]), with more negative amplitudes for right relative to left hemisphere. However, the main effects of Identity (*F*_(1, 24)_ =2.815, *p* = .106, η*_p_^2^* = .105) and Familiarization Stage (*F*_(2, 48)_ = .973, *p* = .385, η*_p_^2^* = .039) were not significant. In addition, all interactions were not significant (all *ps* > 0.05).

**Table 1.** Average (Standard Deviation) of the Peak Amplitudes of the N170 after Three Familiarization Phases.

|  | Left Hemisphere |  |  | Right Hemisphere |  |  |
| --- | --- | --- | --- | --- | --- | --- |
|  | Post-Visual | Post-Identity | Post-Semantic | Post-Visual | Post-Identity | Post-Semantic |
|  | Familiarization | Familiarization | Familiarization | Familiarization | Familiarization | Familiarization |
| Training Identities | -3.98(2.94) | -4.48(3.47) | -4.12(3.30) | -5.50(4.01) | -5.86(3.78) | -5.65(3.76) |
| Control Identities | -3.81(3.15) | -4.17(3.57) | -4.1(3.41) | -5.52(3.79) | -5.62(3.83) | -5.55(3.62) |

**Figure 5.**
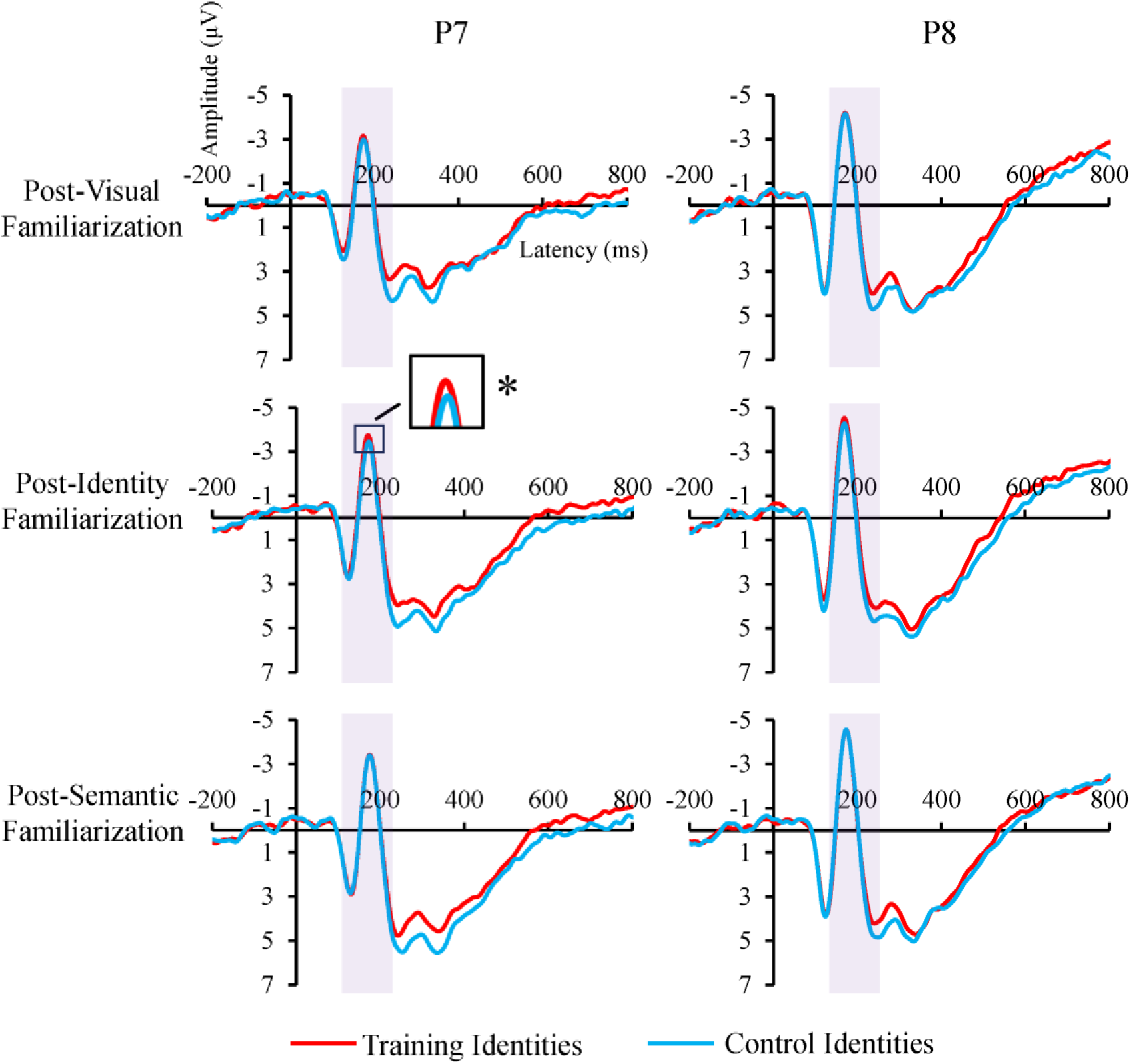
Grand Average Event-Related Potentials of the N170 after Three Familiarization Phases. *Note.* * *p* < .05.

Given that prior studies have linked N170 modulation to identity representation at intermediate stages of familiarization (Caharel & Rossion, 2021), an exploratory repeated-measures ANOVA restricted to the identity familiarization stage was conducted with factors Identity (training, control) and Hemisphere (left, right). This analysis revealed a significant main effect of Identity (*F*_(1, 24)_ =7.826, *p* = .010, η*_p_^2^* = .246, mean difference = .28 μV, 95% CI = [.07, .48]), showing more negative amplitudes for training relative to control identities after identity familiarization. These findings indicate that identity information can modulate the N170 component during face processing. But the main effect of Hemisphere was not significant (*F*_(1, 24)_ = 4.216, *p* = .051, η*_p_^2^* = .149). The interaction of Identity × Hemisphere was not significant (*F*_(1,24)_ = .130, *p* = .722, η*_p_^2^* = .005).

#### 3.2.2. N250

The mean amplitudes of the N250 are shown in Table 2 and Figure 6A & B. The repeated-measures ANOVA revealed a stronger main effect of Identity (*F*_(1, 24)_ = 43.02, *p* < .001, η*_p_^2^* = .642, mean difference = .65 μV, 95% CI = [.45, .86]), with more negative amplitudes for training relative to control identities. The main effect of Familiarization Stage was observed (*F*_(1.52, 36.57)_ = 11.57, *p* < .001, η*_p_^2^* = .325), post-hoc comparison analysis revealed that amplitudes after visual familiarization was more negative amplitudes than after identity familiarization (mean difference = .86 μV, 95% CI = [.38, 1.34]), and semantic familiarization (mean difference = 1.00 μV, 95% CI = [.28, 1.73]). There is no difference between identity familiarization and semantic familiarization (mean difference = .14 μV, 95% CI = [-.65, .36]). However, the main effects of Hemisphere (*F*_(1, 24)_ = .318, *p* = .578, η*_p_^2^* = .013) and all interactions were not significant (all *ps* > 0.05). This indicates that the difference in the N250 between training and control identities emerges after visual familiarization and maintains its stability throughout later phases.

**Table 2.** Average (Standard Deviation) of the Mean Amplitudes of the N250 after Three Familiarization Phases.

|  | Left Hemisphere |  |  | Right Hemisphere |  |  |
| --- | --- | --- | --- | --- | --- | --- |
|  | Post-Visual | Post-Identity | Post-Semantic | Post-Visual | Post-Identity | Post-Semantic |
|  | Familiarization | Familiarization | Familiarization | Familiarization | Familiarization | Familiarization |
| Training Identities | -0.37(2.01) | 0.51(1.93) | 0.67(2.06) | -0.40(2.98) | 0.46(2.44) | 0.38(2.33) |
| Control Identities | 0.23(1.92) | 1.19(2.10) | 1.57(2.12) | 0.20(2.86) | 0.93(2.92) | 1.05(2.40) |

**Figure 6.**
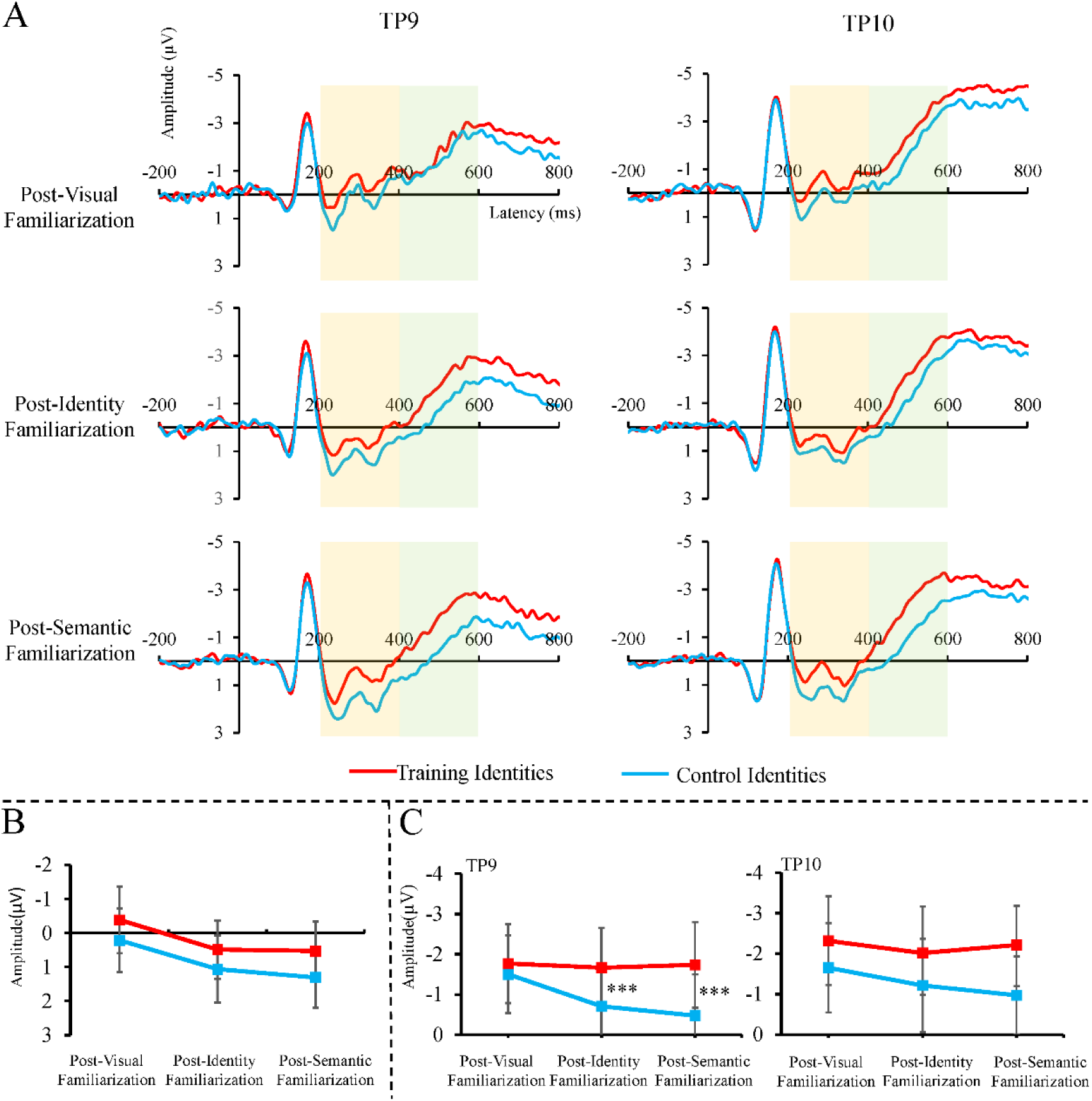
Grand Average Event-Related Potentials and Mean Amplitude Results of N250 and SFE after Three Familiarization Phases. *Note.* A: The waveform of N250 and SFE; B: The results of mean amplitudes of the N250; C: The results of mean amplitudes of the SFE. Error bars for the means indicate 95% CIs. \*\*\**p* < .001

#### 3.2.3. SFE

To capture later, sustained correlates of face familiarity, mean amplitudes in the 400-600 ms time window were analyzed at electrodes TP9 and TP10 (see Table 3 and Figure. 6A & C). The repeated-measures ANOVA revealed a stronger main effect of Identity (*F*_(1, 24)_ = 47.696, *p* < .001, η*_p_^2^* = .665), with more negative amplitudes for training relative to control identities. Neither the main effect of Familiarization Stage (*F*_(2, 48)_ = 1.705, *p* = .193, η*_p_^2^* = .066), nor that of Hemisphere (*F*_(2, 48)_ = 1.208, *p* = .283, η*_p_^2^* = .048) was significant. A significant Identity × Familiarization Stage interaction was observed (*F*_(2, 48)_ = 4.516, *p* = .016, η*_p_^2^* = .158).

**Table 3.** The Average (Standard Deviation) of the Mean Amplitudes of the SFE after Three Familiarization Phases.

|  | Left Hemisphere |  |  | Right Hemisphere |  |  |
| --- | --- | --- | --- | --- | --- | --- |
|  | Post-Visual | Post-Identity | Post-Semantic | Post-Visual | Post-Identity | Post-Semantic |
|  | Familiarization | Familiarization | Familiarization | Familiarization | Familiarization | Familiarization |
| Training Identities | -1.77(2.51) | -1.67(2.52) | -1.74(2.71) | -2.32(2.78) | -2.02(2.65) | -2.22(2.61) |
| Control Identities | -1.50(2.47) | -0.71(2.55) | -0.47(2.62) | -1.65(2.82) | -1.21(2.94) | -0.97(2.46) |

Importantly, a significant three-way interaction among Identity, Familiarization Stage, and Hemisphere was observed (*F*_(2, 48)_ = 3.774, *p* = .030, η*_p_^2^* = .136). To decompose the three-way interaction, separate Identity × Familiarization Stage ANOVA were conducted for each hemisphere. Over the left hemisphere, the main effect of Identity was significant (*F*_(1, 24)_ = 41.63, *p* <.001, η*_p_^2^* = .634), with more negative amplitudes for training identities relative to control identities. The interaction of Identity × Familiarization Stage was significant (*F*_(2, 48)_ = 6.182, *p* = .004, η*_p_^2^* = .205), follow-up comparisons revealed that difference between training and control identities was not significant after visual familiarization (mean difference = .27 μV, 95% CI = [-.75, .22]). However, training identities were elicited more negative amplitudes than control identity after identity familiarization (mean difference = .96 μV, 95% CI = [.50, 1.42]), and after semantic familiarization (mean difference = 1.26 μV, 95% CI = [.91, 1.61]). Over the right hemisphere, only the main effect of Identity was significant (*F*_(1, 24)_ = 41.91, *p* < .001, η*p^2^* = .636, mean difference = .90 μV, 95% CI = [.62, 1.19]), with more negative amplitudes for training relative to control identities across all familiarization stages. Neither the main effect of Familiarization Stage (*F*_(2, 48)_ = 1.152, *p* = .325, η*p^2^* = .046), nor the interaction of Identity × Familiarization Stage (*F*_(2, 48)_ = 2.444, *p* = .098, η*p^2^* = .092) was significant. No other main and interaction effects were significant. These findings indicates that the SFE arises from person-related identity and semantic information rather than visual familiarity alone, and is predominantly expressed in the left hemisphere.

## 4. Discussion

The present study adopted a sequential familiarization paradigm to investigate how distinct types of person-related knowledge (visual exposure, identity information, and semantic knowledge) differentially modulate the three canonical ERP markers of face familiarity. The progressive improvement in recognition performance across the three familiarization stages demonstrates that this sequential paradigm is effective for investigating the cognitive mechanism from face perception to person identification. Importantly, the ERP findings revealed a clear dissociation: the N250 familiarity effect emerged immediately after visual exposure and showed no further enhancement with subsequent familiarization; The N170 exhibited a stage-specific familiarity effect that appeared only after identity familiarization. Most notably, the SFE displayed a clear incremental growth pattern: it was absent after visual familiarization, emerged after identity familiarization, and reached its maximum following the addition of semantic knowledge, particularly in the left hemisphere. Together, these results provide the first direct evidence linking person-related information to specific neural markers of face familiarity, thereby resolving long-standing ambiguities regarding the cognitive origins of these ERP components in familiar face recognition.

The hypothesis that the N250 indexes the accumulation of visual experience was confirmed. Significant N250 amplitude difference between trained and control identities emerged immediately after the visual familiarization, and maintained a stable pattern after the identity and semantic familiarization stages. This finding aligns with prior studies showing that the N250 is modulated by repeated visual exposure (Sommer et al., 2021; Tanaka et al., 2006) and reflects the activation of domain-selective, visual long-term representations of familiar faces (Bojdo et al., 2025; Schweinberger & Neumann, 2016; Wiese et al., 2024). The present results extend these conclusions by demonstrating, within a multi-stage design, that the N250 familiarity effect saturates after perceptual exposure and is insensitive to the later addition of identity or biographical information. This strongly supports the interpretation of the N250 as the electrophysiological correlate of the Face Recognition Unit (Burton et al., 1990; Young & Bruce, 2024), in which a visual memory trace is stably established once sufficient perceptual experience has been accumulated, requiring neither a name nor semantic information (Ghorbani et al., 2025).

In contrast, the N170 showed a stage-specific familiarity effect in the left hemisphere, emerging only after identity familiarization. This finding is consistent with the proposal that the N170 may index early stages of person recognition when robust identity representations have been formed (Caharel & Rossion, 2021). While Caharel and Rossion (2021) concluded from a comprehensive review that the N170 is sensitive to long-term, personally relevant face familiarity, other studies have failed to replicate this effect (Bojdo et al., 2025; Wiese et al., 2019a). Several factors may account for this difference. First, the task employed in the present study was a simple old/new judgment, which does not explicitly require the retrieval of identity-specific information. It is plausible that the N170 familiarity effect is more pronounced when task demands necessitate the active discrimination of individual identities (e.g., Kovács et al., 2023). Second, the intensity and duration of our identity training may not have been adequate to forge the kind of deep, personally relevant representations that typically elicit strong N170 modulations, such as those observed for close friends and relatives (Wiese et al., 2019b). Indeed, Kovács et al. (2023) demonstrated using MVPA that identity-specific information can be decoded from neural activity within the N170 time window. This suggests that identity learning does alter early perceptual processing, but that these changes may manifest as distributed pattern changes rather than robust amplitude shifts in the average ERP. The present univariate N170 effect, though limited, provides crucial behavioral training correlate of this identity-related tuning, highlighting its dependence on the explicit formation of face-name associations.

The most important contribution of the present study concerns the SFE. Our results revealed a clear, stepwise increase in SFE magnitude across the three familiarization stages. This is the first direct demonstration that the SFE is not a single index of face familiarity, but rather a graded neural signal whose strength is parametrically scaled by the depth and richness of the available person-related knowledge. The present findings extend the original characterization of the SFE by Wiese et al. (2019a) in two ways. First, identity information alone was sufficient to initiate the SFE, even without rich semantic or affective content, consistent with evidence that cross-domain identity priming enhances the familiarity response in the 300-400 ms time window (Bojdo et al., 2025). Second, and more importantly, the subsequent addition of biographical knowledge amplified the SFE to its maximal level. This incremental pattern aligns with the notion that the SFE reflects the integration of perceptual face representations with domain-general person knowledge (Wiese et al., 2024; Kovács, 2020), thereby linking the electrophysiological signal to the Person Identity Node (PIN) in classic cognitive models (Burton et al., 1990). The left-hemisphere dominance further implicates the engagement of semantic processing systems, consistent with the hub-and-spoke framework in which anterior temporal regions serve as a convergence zone for conceptual knowledge (Patterson & Ralph, 2016; Ralph, 2014). The SFE may thus represent a re-entrant feedback signal that broadcasts the successful retrieval of person-specific conceptual information back to perceptual face-processing regions (Kovács, 2020).

## 5. Conclusion

In sum, by employing a sequential familiarization paradigm, this study provides the first direct evidence that the N250, N170, and SFE are driven by distinct types of person-related information: visual experience, identity information, and semantic knowledge, respectively. These findings reveal the information-specific origins of the core ERP components of face familiarity, and they illuminate the electrophysiological trajectory through which the accumulation of person-related knowledge transforms a perceptual face representation into the recognition of a known person.

## Notes

### Competing Interest Statement

The authors have declared no competing interest.

